# Hox genes are essential for the development of novel serial homologous eyespots on the wings of *Bicyclus anynana* butterflies

**DOI:** 10.1101/814848

**Authors:** Yuji Matsuoka, Antónia Monteiro

**Affiliations:** Department of Biological Sciences, National University of Singapore, 14 Sciences Drive 4, 117543 Singapore; Science Division, Yale-NUS College, 10 College Avenue West, 138609 Singapore

## Abstract

The eyespot patterns found on the wings of nymphalid butterflies are novel serial homologous traits that originated first in hindwings and subsequently in forewings, suggesting that eyespot development might be dependent on Hox genes. Hindwings differ from forewings in the expression of *Ultrabithorax* (*Ubx*), but the function of this Hox gene in eyespot development as well as that of another Hox gene *Antennapedia* (*Antp*), expressed specifically in eyespots centers on both wings, are still unclear. We used CRISPR-Cas9 to target both genes in *Bicyclus anynana* butterflies. We show that *Antp* is essential for eyespot development on the forewings and for the differentiation of white centers and larger eyespots on hindwings, whereas *Ubx* is essential for the development of at least some hindwing eyespots but also for repressing the size of other eyespots. Additionally, *Antp* is essential for the development of silver scales in male wings. In summary, *Antp* and *Ubx*, in addition to their conserved roles in modifying serial homologous traits along the anterior-posterior axis of animals, have acquired a novel role in promoting the development of a new set of serial homologs, the eyespot patterns, in both forewings (*Antp*) and hindwings (*Antp* and *Ubx*) of *B. anynana* butterflies. We propose that the peculiar pattern of eyespot origins on hindwings first, followed by forewings, could be due to an initial co-option of *Ubx* into eyespot development, followed by a later, partially redundant co-option of *Antp* into the same network.

## Introduction

Hox genes are primarily known for their embryonic expression domains of broad stripes along the anterior-posterior axis of bilaterian animals and for giving body regions along this axis a unique identity (Lewis 1978, McIntyre et al., 2007, Mallo et al., 2010). In arthropod animals, these unique identities are often visualized by changes in the external appearance of serial homologous traits along the body, such as appendages. However, Hox genes have not been implicated in the origin of appendages, but rather in their modification or repression by tweaking the appendage’s gene regulatory networks (GRN) (Slattery et al., 2011). This is because the silencing of Hox genes changes an appendage’s identity, or leads to the development of additional appendages, rather than cause the loss of the appendage itself (Struhl 1982, Carroll et al., 1995, Tomoyasu et al. 2005; Ohde et al. 2013; Martin et al. 2015).

Aside from their classic role as modifiers of serial homologs, Hox genes have also been secondarily recruited to contribute to the development of lineage-specific evolutionary novelties. Here, Hox genes have acquired novel expression domains, often post-embryonically, and appear to contribute to the development of a novel trait. For example, a novel *Sex combs reduced* (*Scr*) expression domain in the first pair of pupal legs of flies is required to produce sex combs used for mating (Tanaka et al. 2011). A late *Scr*’s pre-pupal and pupal expression in pronotal horns in beetles regulates horn development (Wasik et al. 2010). A novel expression domain of *Ubx* in the tibia of third thoracic embryonic legs leads to the development of the pollen basket in worker bees (Medved et al., 2014). And, the late pupal expression of *Abd-B* in *Drosophila* and *Bombus* abdominal epidermis leads, in both cases, to the origin of novel pigmentation patterns in the abdomen of these insects (Jeong et al., 2006, Tian et al., 2019). In these examples, and quite distinctly from their function as modifiers of serial homologs, when a Hox gene is disrupted, the novel trait is severely disrupted or lost.

Eyespots on the wings of nymphalid butterflies are an interesting example of both a novel trait, as well as a serial homologous trait, but the molecular changes that led to the origin of these traits are still unknown. Ancestral state reconstructions on a large phylogeny of ∼400 genera suggested that eyespots first originated in four to five wing sectors on the ventral side of the hindwings of an ancestral lineage of nymphalid butterflies before appearing on forewings or dorsal sides of both wings (Oliver et al. 2014; Schachat et al., 2015). The origin of eyespots restricted to hindwings is intriguing and could suggest that hindwing-specific Hox genes might have been required for eyespot origins.

Butterflies have up to two known candidate Hox genes, *Ubx* and *Antp*, expressed on their larval hindwings (Fig. 1A, Weatherbee et al. 1999; Saenko et al., 2011). *Ubx* is expressed homogeneously across the whole hindwing, as observed in most other insects (Prasad et al., 2016), whereas *Antp* has a novel and more specific expression pattern in the eyespot centers in both the fore- and hindwing disks in only a subset of butterflies with eyespots (Saenko et al., 2011; Shirai et al., 2012; Oliver et al., 2012, Tong et al., 2014). Interestingly, in *Bicyclus anynana, Ubx* is additionally expressed at slightly elevated levels in the future eyespot centers in larval and pupal hindwings (Fig. 1A, Tong et al. 2014), whereas *Junonia coenia* does not express *Antp* nor elevated *Ubx* expression in their eyespots (Weatherbee et al. 1999; Tong et al. 2014; Shirai et al., 2012; Oliver et al., 2012). These observations suggest that *Ubx* homogeneous hindwing expression alone, but not *Antp* nor *Ubx* eyespot-specific expression, might have played some role in eyespot origins. However, *Ubx* cannot be responsible for the origin of eyespots on forewings since it is not expressed there in any of the species examined so far, unlike *Antp*.

**Fig 1.**
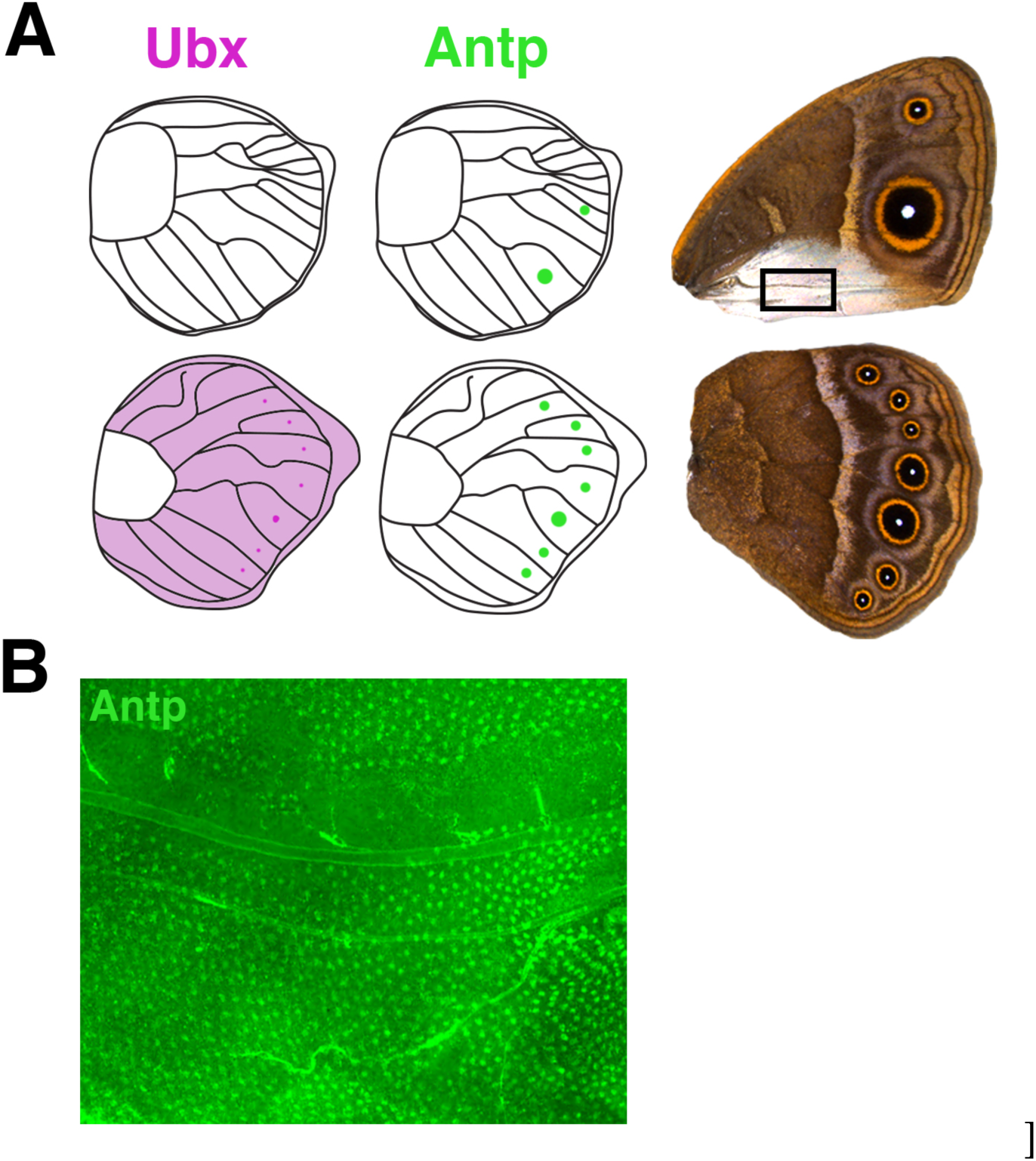
Expression pattern of Ubx and Antp in wings. (A) Schematic diagram of the expression pattern of Ubx and Antp on the larval wings and corresponding adult male wings. Ubx is expressed homogeneously across the hindwings and more intensely in the eyespot centers. Antp is expressed in all eyespot centers. (B) Expression of Antp in a section of a Wt male pupal forewing corresponding to the rectangle in (A). Antp is expressed intensely in silver scale building cells (arrowheads) during the pupal stage (Image courtesy of Xiaoling Tong).

To date, neither *Ubx* nor *Antp* have been directly targeted for loss-of-function experiments in any butterfly species, and the role of these genes in eyespot development remains unclear. Here, we investigate the functions of *Antp* and *Ubx* in eyespot development in *B. anynana* with CRISPR-Cas9 to create mosaic mutants. We show that *Antp* is required for eyespots to form on the forewings, whereas *Ubx* has both a repressive and activating role on eyespot formation on the hindwings, depending on wing position. By integrating these results with previous work detailing the origin of eyespots across wings and wing surfaces of nymphalid butterflies, we propose that Hox genes were probably implicated in the origin of these novel traits in hindwings (first) and forewings (second) during butterfly diversification.

## Results

### *Antp* crispants lose forewing eyespots and produce smaller hindwing eyespots with no white center

*Antp* crispant butterflies had distinct phenotypes on fore- and hindwings. On the forewings, *Antp* crispants lost both the anterior (M1) and posterior (Cu1) eyespots on both ventral and dorsal surfaces (Compare Fig. 2A with 2B and 2C). This was also observed in on dorsal surfaces (Fig. 2J). Furthermore, the silver scales observed at the posterior end of the male ventral forewings were transformed to brown background scales, which are observed at this location only in females (Figs 2C and 2D). In male *B. anynana*, the expression of *Antp* in the early pupal stage was associated with the development of silvery scales on the forewings (Fig. 1B and Fig. 2C), which was a previously undocumented expression pattern for this gene. On the hindwings of *Antp* crispants, the white eyespot centers were missing in every eyespot (on both ventral and dorsal surfaces) (Fig. 2F, 2H and 2K), and the overall size of the eyespots was reduced (Fig. 2H, 2I compared to 2G), but the eyespots were never lost (Fig. 2H). These results indicate that *Antp* is essential for eyespot development and silver scale development in forewings, whereas in hindwings, *Antp* seems to be only required for the differentiation of the white scales and for increasing the size of the eyespots.

**Fig 2.**
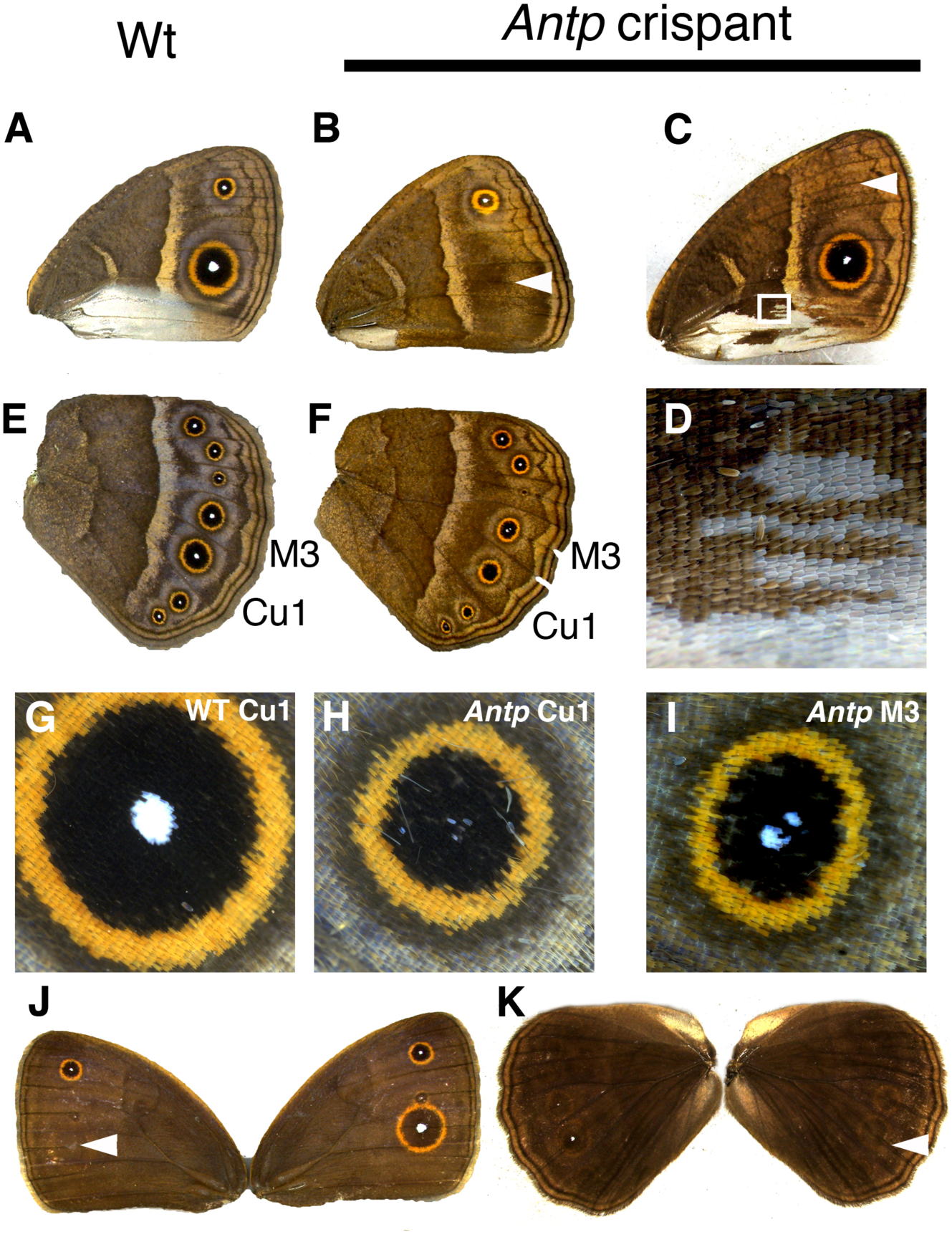
*Antp* crispant phenotypes on the adult wings. Ventral side of a Wt male forewing. (B and C) Ventral side of *Antp* crispant forewings. (B) *Antp* crispants partially or completely lost the posterior eyespot, or lost the (C) anterior eyespot (marked with white arrowheads). Male-specific silver scales on the posterior part of the forewing were partially transformed into brown background scales (close up in D). (E) Ventral side of Wt male hindwing. (F) Ventral side of *Antp* crispant male hindwing. *Antp* crispants partially or completely lost white eyespot centers, but eyespots were never missing. (G) Cu1 eyespot from Wt hindwing. (H and I) Highly magnified pictures of eyespots from F. The white eyespot center was completely or partially lost, and the eyespots were reduced in size. (J and K) Dorsal side of *Antp* crispant fore- and hindwings. Eyespots were also lost on the dorsal side for both fore- and hindwings (white arrowheads).

### *Ubx* crispants led to the homeotic transformation of hindwings into forewings by both the activation and repression of eyespots

*Ubx* crispants showed the predicted homeotic phenotype, e.g., of hindwings acquiring a forewing identity (Fig. 3A). In particular, hindwing M3 eyespots disappeared (Fig. 3B compared to 3C), and hindwing Cu1 eyespots became as large as the corresponding Cu1 eyespots on the forewing (Fig 3D compared to 3E). In addition, male-specific silver scales, which are normally present on the posterior edge of the forewing, developed ectopically at a homologous posterior position in hindwings of crispant males (Fig 3F compared to 3G). These results indicate that *Ubx* has both repressive as well as activating effects on eyespots, depending on their position on the wing, e.g., *Ubx* has a repressive role on Cu1 eyespot size, and an activating, essential role on M3 eyespot development. In addition, *Ubx* is a repressor of silver scale development on posterior ventral hindwings.

**Fig 3.**
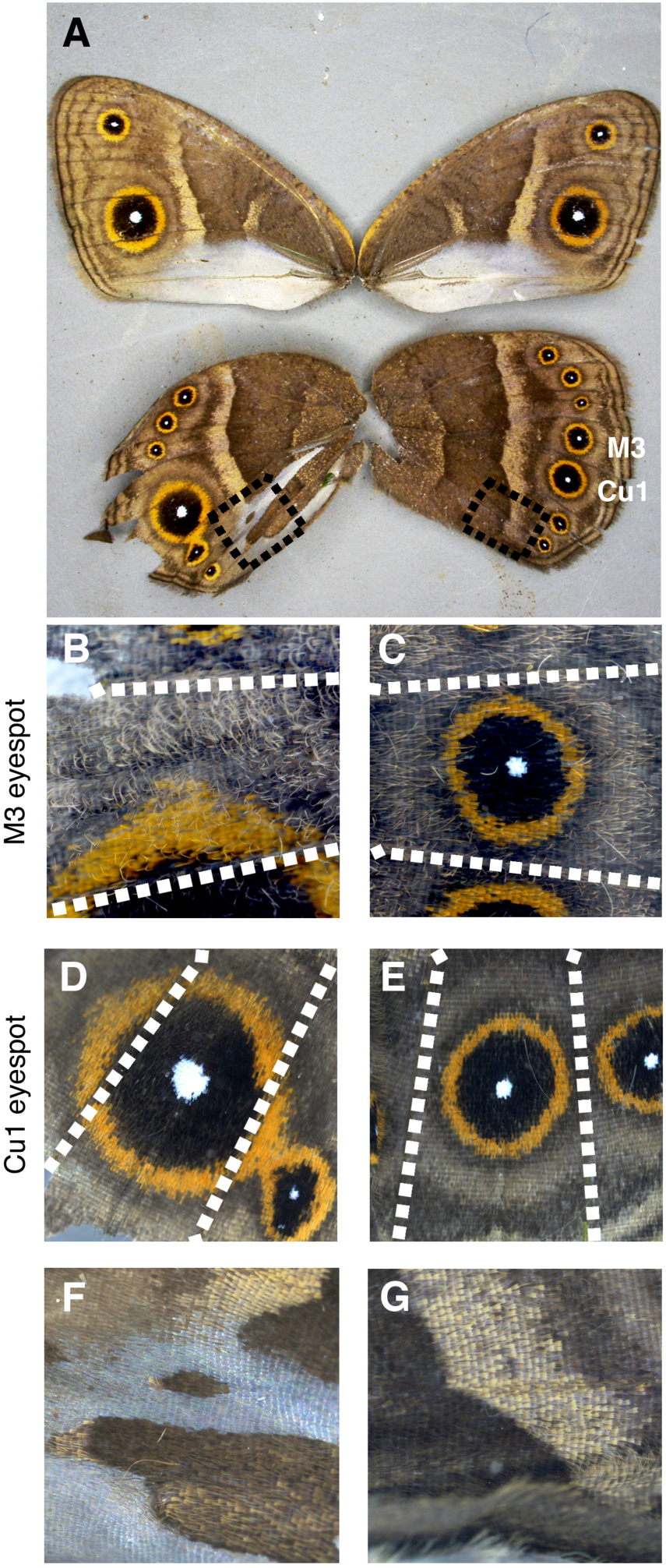
A *Ubx* crispant individual (ventral side). The hindwing on the left is highly mutated, whereas the wings on the right are not and represent the wildtype wing pattern. (B) The M3 eyespot from the *Ubx* crispant is missing, resembling the forewing, while it is present on the other wing (C). White dotted lines indicate the position of veins bordering the M3 sector. (D) The Cu1 hindwing eyespot became enlarged, resembling the forewing Cu1 eyespot, relative to the non-mutated Cu1 eyespot on the wing shown in (E). (F, G) Enlargements of the black dotted areas in A. (F). Ectopic silver scales were generated on the posterior hindwing of the *Ubx* crispant instead of brown scales normally found at this position in non-mutated wings (G).

## Discussion

Here we show that the Hox genes *Antp* and *Ubx* have evolved novel expression domains and functions in *B. anynana* beyond those connected to anterior-posterior axis patterning of embryos that give segments and appendages along this axis their unique identities. Our results implicate these genes in the development of silver scales, and in the development of a morphological innovation within the Lepidoptera, eyespots, pointing to their likely involvement in eyespot origins.

### *Antp* is essential for eyespot development on forewings only

*Antp* had been hypothesized to function in eyespot development in fore- and hindwings because of its expression pattern in the eyespot centers of both larval (Fig.1A, Saenko et al. 2011; Shirai et al. 2012; Oliver et al. 2012) and pupal wings (Tong et al. 2014). Here, we provide the first functional evidence supporting this hypothesis: *Antp* is an essential gene for eyespot development in forewings. In hindwings, however, *Antp* is only required for the differentiation of the white eyespot centers and for an increase in eyespot size. The different functions of *Antp* in fore- and hindwing eyespots may be related to the isolated expression of *Antp* in forewing eyespots, and the co-expression of *Antp* and *Ubx* in hindwing eyespots. On the forewings, *Antp* is likely to be required for both focal establishment (in the larval stage) and the production of the morphogenetic signal in early pupal stages that differentiates the color rings. It has been suggested that the white scales at the eyespot centers may need high levels, the black scales moderate levels, and golden scales lower levels of a morphogenetic signal to differentiate into their respective colors (French and Brakefield 1995, Brakefield et al. 1996). On the hindwings, however, *Ubx* might be able to partially substitute for these roles of *Antp*. When *Antp* activity is removed, eyespot foci are still able to differentiate but might not be able to generate enough signal to differentiate the central white scales, nor to reach the same number of cells away from the center, leading to overall smaller eyespots.

### *Ubx* acts both as a repressor and as an essential gene for eyespot development

In this study, we showed that hindwing Cu1 eyespots became larger in *Ubx* crispants (Fig. 3D), while M3 eyespots disappeared (Fig. 3B), suggesting opposite and location-specific effects of *Ubx* on eyespot development. The enlargement of Cu1 eyespots is consistent with a previously proposed repressor function for *Ubx* on both *J. coenia* and *B. anynana* Cu1 eyespots (Weatherbee et al. 1999, Tong et al. 2014). These prior experiments made use of a spontaneous mutant line of *J. coenia* that developed patches of forewing color patterns on the hindwing and also lacked *Ubx* expression in clones of hindwing cells (the nature of the mutation still remains to be characterized). When clones of transformed cells included Cu1 eyespots, these eyespots were transformed into larger eyespots bearing the size and colors of forewing eyespots. Furthermore, in *B. anynana*, the overexpression of *Ubx* caused a reduction of Cu1 eyespot size in both fore- and hindwings, again suggesting a repressor function of *Ubx* on Cu1 eyespot size regulation (Tong et al., 2014). However, the disappearance of M3 hindwing eyespots in *Ubx* crispants clearly supports a novel, previously undocumented, eyespot-promoting function for this gene.

While we have no clear insight for why *Ubx* acts in such different ways towards eyespots in different sectors of the wing, we speculate that these different modes of action might be accomplished in two possible ways: 1) by the Ubx protein having distinct types of downstream targets: different sector-specific selector genes that are present in only some sectors of the wing (e.g., *engrailed, invected, spalt, optomotor blind*, or their downstream targets) (Carroll et al., 1994, Keys et al., 1999; Monteiro 2015; Özsu et al., 2017) and which, in turn, interact with the eyespot GRN by activating it or repressing it, or 2) by the Ubx protein using these sector-specific selectors as cofactors to either activate or repress the eyespot GRN in different ways in the different sectors (Mann et al., 2009). In summary, we propose that *Ubx* functions both as an activator and a repressor of eyespots in species, such as *Bicyclus* and *Junonia*, via its interactions with sector-specific selector genes.

### *Antp* promotes, while *Ubx* represses silver scale development

Silver scales develop only in males on the ventral posterior side of forewings and on the dorsal anterior side of hindwings, closely associated with scent glands that synthesize and release male sex pheromones (Fig. 1C) (Dion et al., 2016). In *Antp* crispants, patches of silver scales on the forewing were changed to brown scales (Fig. 2C and 2D), whereas in *Ubx* crispants, ectopic silver scales were generated at the posterior end of the hindwing, which are normally covered by brown scales (Fig. 3F). These results suggest that *Antp* promotes, whereas *Ubx* represses silver scale development. Our lab recently showed that the male isoform of the sex determination pathway gene, *doublesex* (*dsx*), is also required for silver scale development in *B. anynana* males on both fore- and hindwings (Prakash and Monteiro 2019). In addition, the dorsal selector gene *apterous A* (*apA*) regulates silver scale development in a surface-specific manner in males: it represses silver scale development from dorsal forewings and promotes silver scale development on dorsal hindwings (Prakash and Monteiro 2018). We speculate that the male isoform of *dsx* activates *Antp* expression at the posterior end of the forewing to produce silver scales. *Ubx* might repress the expression of *Antp, dsx*, or any of their downstream targets, at the posterior end of the hindwing to prevent the generation of silver scales at the homologous hindwing region.

### Possible functions of *Antp* and *Ubx* in eyespot evolution

*Antp* and *Ubx* phenotypes give us additional insights about the evolution of eyespot number and location on the wings of nymphalid butterflies. Here we propose that since forewings and hindwings differ in the expression of a key Hox gene, *Ubx*, which was shown here to be required to activate eyespot deployment (in M3 sectors), this gene might have been essential for the origin of eyespots, which was limited to the hindwings (Fig. 4) (Oliver et al. 2014). Recent work has identified a reaction-diffusion (RD) mechanism involving the gene *Distal-less* (*Dll*) in promoting the differentiation of eyespot centers in larval wings (Connahs et al. 2019). If novel binding sites for Ubx (or for a Ubx cofactor or target) evolved in the regulatory regions of any of the genes involved in this RD mechanism, and if this led to the stabilization of *Dll* expression in eyespot centers, this might have aided the origin of eyespots in hindwings only. Eyespots appear to have subsequently evolved on forewings multiple times independently in different lineages (Schachat et al, 2015). We propose that *Antp* was required for eyespots to eventually originate on the forewings of butterflies from the subfamilies *Satyrinae* and *Biblidinae*. The co-option of *Antp* to the eyespot gene regulatory network (GRN) in these lineages may have allowed eyespots to be activated, for the first time on the forewing, as this Hox gene might have substituted for the activating role of *Ubx* on this novel wing surface, and also led to an increase in the overall size of hindwing eyespots. The recruitment of *Antp* to the eyespot GRN also led to the origin of the white centers (Fig. 4), at least in the lineage leading to *B. anynana*. Genes other than *Antp* might have allowed eyespots to emerge in forewings in other lineages of nymphalids that do not express *Antp* in eyespots, such as *Junonia*. Furthermore, negative regulators of the eyespot GRN that are expressed exclusively on dorsal wing surfaces, such as *apA*, might have further limited the origin of eyespots to the ventral surfaces of wings (Prakash and Monteiro 2018). Finally, eyespots appear to have originated on the dorsal surfaces via the repression of *apA* (via a yet unidentified mechanism), as observed in the region of the anterior (M1) and posterior (Cu1) eyespot centers in *B. anynana* (Prakash and Monteiro 2018). Variation in eyespot size and number between forewings and hindwings has likely further evolved via novel interactions between the Hox genes and the sector-specific selector genes that still need to be identified. Further comparative investigations will be needed to test whether the currently known role of *Ubx* as a repressor of Cu1 eyespot size in two divergent lineages of nymphalids (*Bicyclus* and *Junonia*) represents the ancestral function of this gene, or if this is a more derived function that evolved separately in each lineage. It is worth noting that close relatives of each of these species can vary dramatically in the relative size of their Cu1 forewing and hindwing eyespots, arguing for two separate origins for this function in *Ubx*.

**Fig 4.**
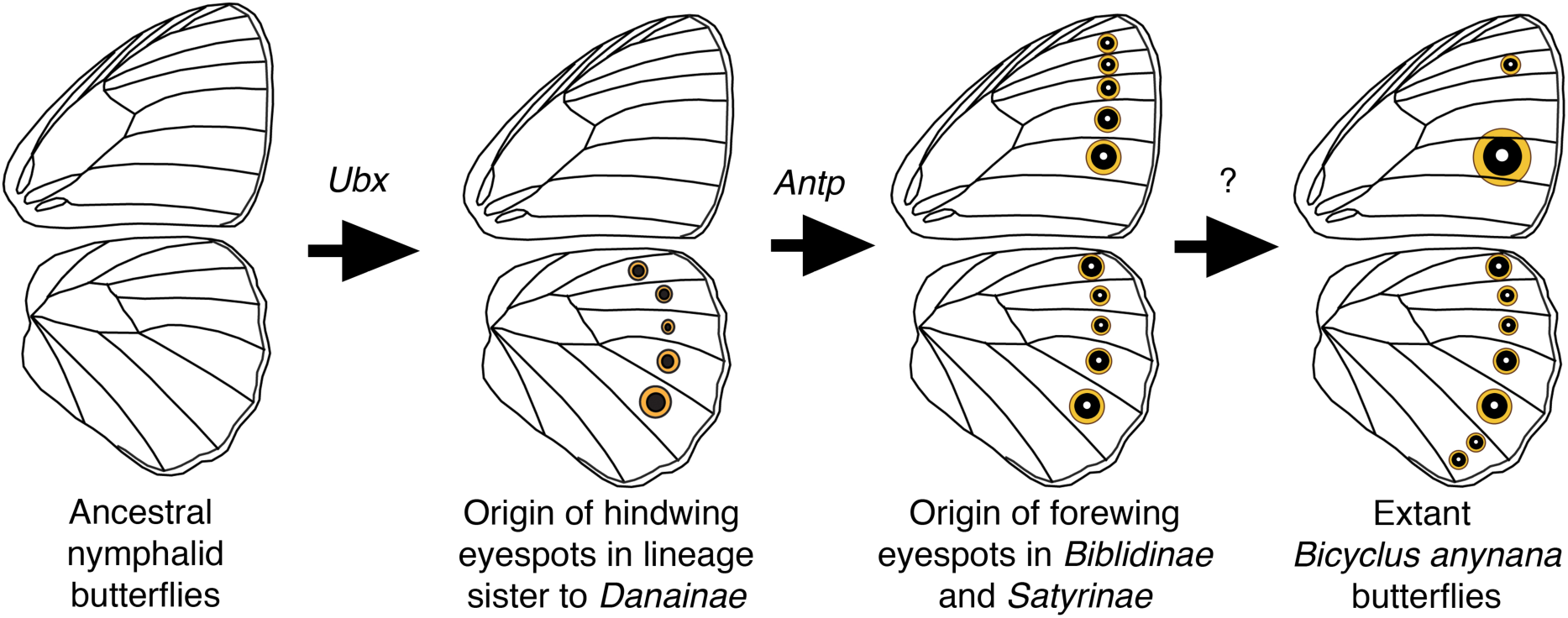
Possible functions of *Ubx* and *Antp* in eyespot origins. Ancestral state reconstructions suggested that eyespots first originated in four to five wing sectors on the ventral side of the hindwings. Eyespots later appeared on the ventral sides of the forewing (and later on the dorsal sides of both wings). We propose that *Ubx*, shown here to be essential for the activation of some hindwing eyespots, was instrumental in the origin of eyespots restricted to hindwings. Once *Antp* was co-opted to the eyespot GRN, its functional similarity to *Ubx* permitted eyespots to develop on the forewings (in lineages of butterfly that express this gene in eyespot centers), and also to become larger (in hindwings) as well as acquire a white center.

In conclusion, we report novel functions for the Hox genes *Antp* and *Ubx* that implicate these genes in the development and origin of nymphalid eyespots, a new system of serial homologs. Our data also shed light on the mechanisms that led to the evolution of differences in eyespot number, size, and morphology between fore- and hindwings. In particular, we propose a novel hypothesis for eyespot origins: that *Ubx* was essential in restricting the origin of eyespot serial homologs to hindwings, and that the recruitment of *Antp* to the eyespot GRN led to redundancy of function (with *Ubx*) and to the appearance of eyespots on forewings, at least in the satyrid lineage of butterflies. Finally, our work also implicates *Antp* in the development of white eyespot centers and silver scales in *B. anynana*. Future comparative work should examine how the function and deployment of these genes has evolved at a finer scale across a butterfly phylogeny to test the eyespot origin hypothesis further.

## Materials and methods

### Butterfly husbandry

*B. anynana*, originally collected in Malawi, have been reared in the lab since 1988. The caterpillars were fed on young corn plans and adults on mashed banana. *B. anynana* were reared at 27°C and 60% humidity in a 12:12 light:dark cycle.

### sgRNA design

sgRNA target sequences were selected based on their GC content (around 60%) and the number of mismatch sequences relative to other sequences in the genome (> 3 sites). In addition, we selected target sequences that started with a guanidine for subsequent *in vitro* transcription by T7 RNA polymerase.

### sgRNA production

The template for *in vitro* transcription of sgRNA was made with a PCR method described in Matsuoka and Monteiro (2018) and Banerjee and Monteiro (2018; containing a video protocol). The forward primer contains a T7 RNA polymerase binding site and a sgRNA target site (GAAATTAATACGACTCACTATAGNN_19_GTTTTAGAGCTAGAAATAGC). The reverse primer contains the remainder of sgRNA sequence (AAAAGCACCGACTCGGTGCCACTTTTTCAAGTTGATAACGGACTAGCCTTATTTTAACT TGCTATTTCTAGCTCTAAAAC). PCR was performed with Q5 High-Fidelity DNA Polymerase (NEB) in 100 µL reaction volumes. After checking with gel electrophoresis, the PCR product was purified with the Gene JET PCR purification kit (Thermo Fisher). *In vitro* transcription was performed with T7 RNA polymerase (NEB), using 500 ng of purified PCR product as a template for overnight. After DNase I treatment to remove the template DNA with, the RNA was precipitated with ethanol. The RNA was then suspended in RNase-free water and stored at −80□.

### Cas9 mRNA production

The plasmid pT3TS-nCas9n (Addgene) was linearized with Xba□ (NEB) and purified by phenol/chloroform purification and ethanol precipitation. *In vitro* transcription of mRNA was performed using the mMESSAGEmMACHINE T3 kit (Ambion). One μg of linearized plasmid was used as a template, and a poly(A) tail was added to the synthesized mRNA by using the Poly(A) Tailing Kit (Thermo Fisher). The A-tailed RNA was purified by lithium-chloride precipitation and then dissolved to RNase-free water and stored at −80□.

### Microinjection

Eggs were laid on corn leaves for 30 min. Within 2-3 h after egg laying, sgRNA and Cas9 mRNA were co-injected into embryos. The concentrations of sgRNA and Cas9 are listed in Table 1. Food dye was added to the injection solution for better visualization. The injections were performed while the eggs were submerged in PBS. The injected eggs were incubated at 27□ in PBS, transferred onto moist cotton on the next day, and further incubated at 27□. The hatched caterpillars were moved to corn leaves and reared at 27□ with a 12:12 h light:dark cycle and 60% relative humidity.

**Table 1.** Summary of injection.

| Target | sgRNA final concentration | Cas9 mRNA final Concentration | No. of injected embryos | No. of hatched larvae | No. of adults | No. of adults showing phenotype |
| --- | --- | --- | --- | --- | --- | --- |
| *Antp* #1 | 0.5 µg/µl | 1 µg/µl | 176 | 61 | 18 | 1 |
| *Antp* #1 | 0.1 µg/µl | 0.2 µg/µl | 111 | 47 | 28 | 1 |
| *Antp* #1 | 0.1 µg/µl | 0.2 µg/µl | 146 | 59 | 10 | 1 |
| *Antp* #2 | 0.1 µg/µl | 0.2 µg/µl | 161 | 38 | 7 | 3 |
| *Ubx* #1, 2 | 0.4 µg/µl each | 0.8 µg/µl | 151 | 59 | 21 | 1 |
| *Ubx* #1, 2 | 0.3 µg/µl each | 0.6 µg/µl | 142 | 17 | 5 | 0 |
| *Ubx* #1, 2 | 0.25 µg/µl each | 0.5 µg/µl | 182 | 46 | 22 | 1 |

### Detection of indel mutations

Genomic DNA was extracted with the SDS and Proteinase K method from a pool of 5 injected embryos that did not hatch. About 250 bp of sequence spanning the target sequence was amplified with PCRBIO Taq Mix Red (PCRBIOSYSTEMS), and PCR conditions were optimized until there was no smear, primer dimers or extra bands. Primers for those analyses are listed in Table 2. The PCR products were purified with the Gene JET PCR purification kit (Thermo Fisher). Two hundred nanograms of PCR product were denatured and re-annealed in 10x NEB2 buffer. One microliter of T7 endonuclease □ (NEB) was added to the sample, while 1 µl of MQ water was added to a negative control. Immediately after the incubation for 15 min at 37□, all the reactions were analyzed on a 3% agarose gel. Amplicons that showed positive cleavage from the T7 endonuclease □ assay were subcloned into the pGEM-Teasy vector (Promega) through TA cloning. For each target, we picked 8 colonies and extracted the plasmid with a traditional alkali-SDS method and performed a PEG precipitation. Sequence analysis was performed with the BIGDYE terminator kit and a 3730*xl* DNA Analyzer (Thermo Fisher).

**Table 2.**
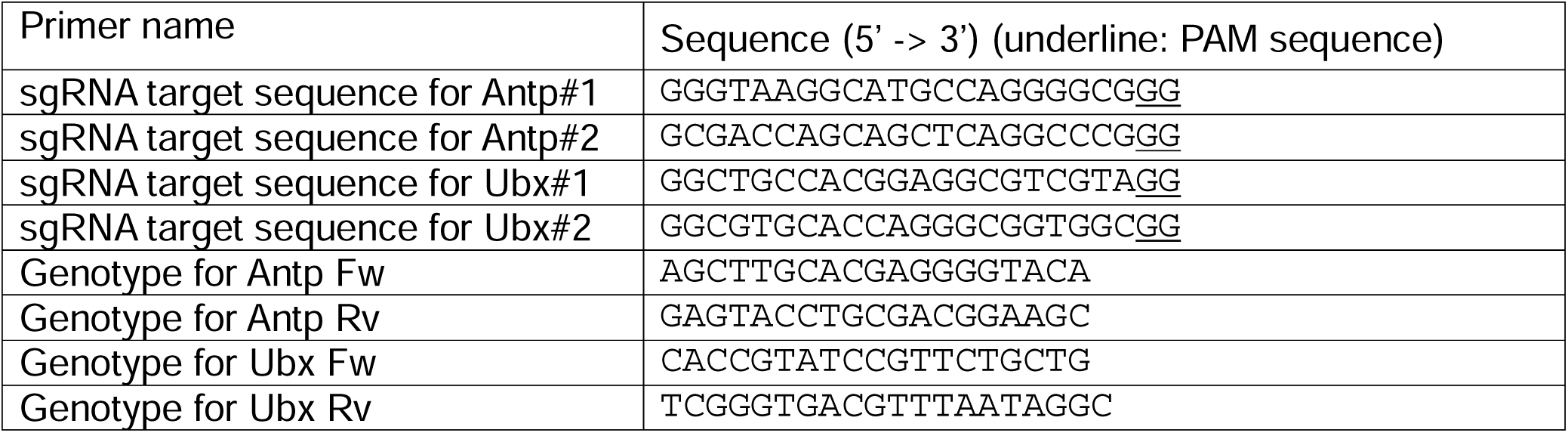
Primer list.

## Acknowledgements

We thank Xiaoling Tong for providing a high-resolution image of a pupal wing stained with Antp, and Thomas Werner and Anupama Prakash for comments on the manuscript. Funding was from grant MOE2015-T2-2-159 from Ministry of Education, Singapore, and from the National Research Foundation, Singapore award NRF-NRFI05-2019-0006.

---

**Fig S1.**
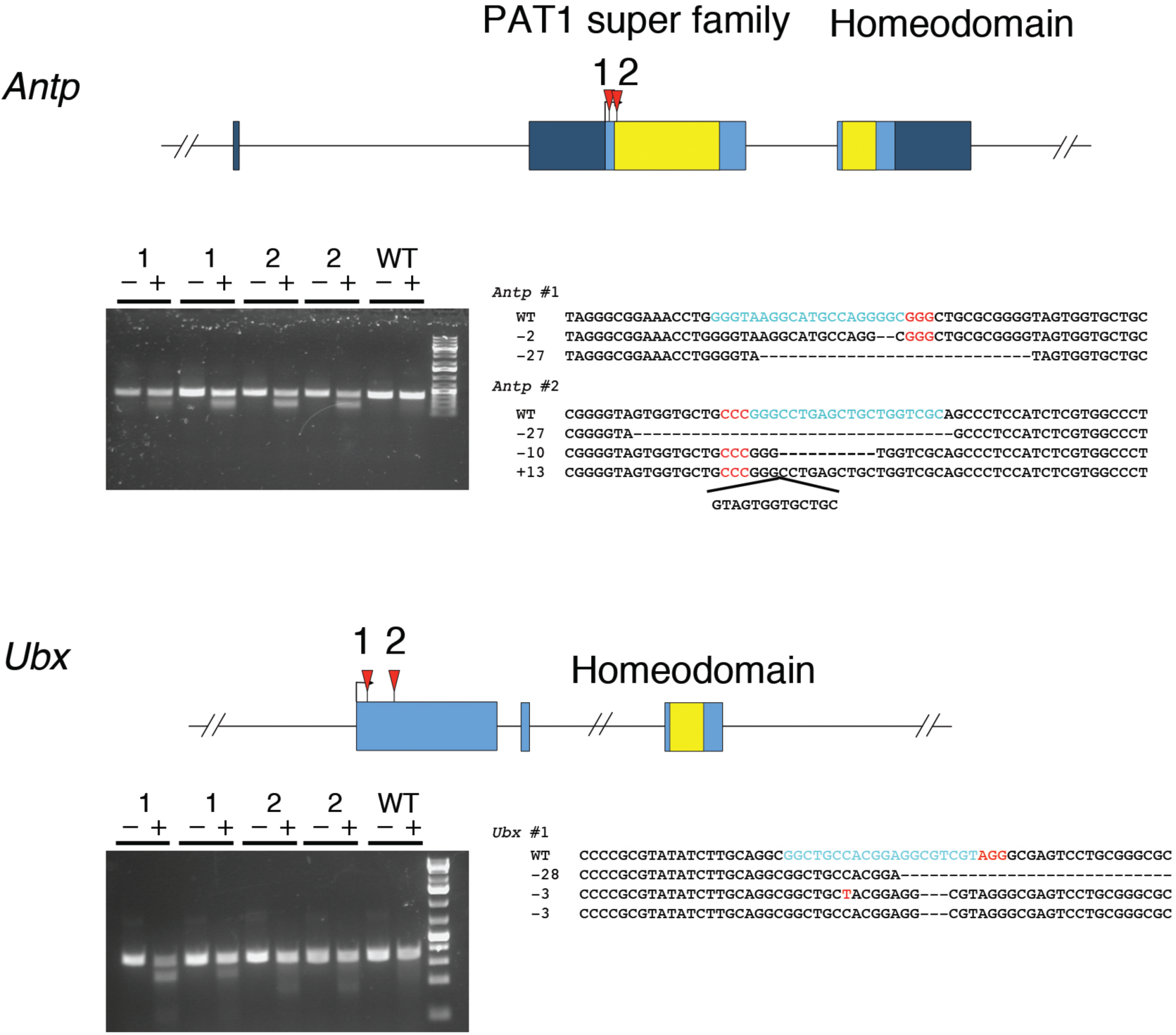
T7 endonuclease I assay and sequence analysis for CRISPR/Cas9 mutations. Schematic representation of gene structure of target genes. Blue boxes indicate exons, black v-shaped lines connecting boxes indicate introns, and dark blue boxes indicate untranslated regions. Yellow-colored regions inside exons indicate functional domains. Each functional domain was annotated using a conserved domain search at NCBI. Red arrowheads indicate the CRISPR target region. The gel shows the result of a T7 endonuclease assay performed on embryos after injection of guide and Cas9. “Minus” lanes indicate a negative control where T7 was not added to the reaction. “Plus” lanes indicate the presence of T7 enzyme. The expected sizes of digested bands were observed only from the lanes having T7 enzyme. Sanger sequence results indicate that an indel mutation was generated around the target site.

